# Loss of the E3 ubiquitin ligase TRIM67 alters the post-synaptic density proteome

**DOI:** 10.1101/2024.01.05.574385

**Authors:** Laura E. McCormick, Natalie K. Barker, Laura E. Herring, Stephanie L. Gupton

## Abstract

The E3 ubiquitin ligase TRIM67 is enriched in the central nervous system and is required for proper neuronal development. Previously we demonstrated TRIM67 coordinates with the closely related E3 ubiquitin ligase TRIM9 to regulate cytoskeletal dynamics downstream of the netrin-1 during axon guidance and axon branching in early neuronal morphogenesis. Interestingly, loss of *Trim67* impacts cognitive flexibility in a spatial learning and memory task. Despite this behavioral phenotype, it was previously uninvestigated if TRIM67 was involved in synapse formation or function. Here we demonstrate TRIM67 localizes to the post-synaptic density (PSD) within dendritic spines. Furthermore, we show that loss of *Trim67* significantly changes the PSD proteome, including changes in the regulation of the actin and microtubule cytoskeletons. Collectively, our data propose a synaptic role for TRIM67.

## Description

Ubiquitination is a post-translational modification that influences protein degradation, localization, and function. E3 ubiquitin ligases add the 8 kDa ubiquitin protein onto a substrate. There are more than 600 mammalian E3 ligases encoded in the genome; research unveiling novel functions of each ligase and their substrate specificity across cell types is ongoing. The brain-enriched class I TRIM ligase TRIM67 has been implicated in neuronal development, particularly in early stages of neuronal shape change. For example, knockdown of TRIM67 in immortalized neuroblastoma cells reduces neurite formation (Yaguchi et al. 2012). Furthermore, cortical neurons from *Trim67*^*-/-*^ mice have shorter axons that do not respond to the axon guidance cue netrin-1 and display dysregulated dynamics of the actin cytoskeleton (Boyer et al. 2020). The *Trim67*^-/-^ mouse displays brain hypotrophy, as well as reduced axon fiber thickness in the corpus callosum and hippocampus (Boyer et al. 2018). Lastly, loss of murine *Trim67* results in several behavioral phenotypes, including reduced cognitive flexibility in the Morris Water Maze test (Boyer et al. 2018), an experiment designed to measure spatial memory and learning.

Recently, we demonstrated that TRIM9, the gene paralog of TRIM67, localizes to the synapse and is required for netrin- 1-dependent signaling in dendritic spines. Furthermore, loss of *Trim9* altered synaptic protein levels in vitro and in vivo (McCormick et al. 2024). Notably, TRIM9 and TRIM67 are independently required for morphogenetic responses to netrin-1 during axon pathfinding and share a multitude of interaction partners (Boyer et al. 2020; Menon et al. 2021). These similar, but non-redundant functions, combined with the behavioral phenotypes of *Trim67*^-/-^ mice (Boyer et al. 2018) led us to hypothesize TRIM67 may also localize to the synapse and influence synaptic function. To determine if TRIM67 localizes to the pre- or post-synapse, we utilized differential centrifugation of murine forebrains to enrich for the synaptosome fraction. The synaptosome is further separated into the presynaptic and post-synaptic density (PSD) enriched fractions (**Fig 1A**). The PSD is a specific component of dendritic spines containing a multitude of proteins essential for neurotransmitter reception and the concomitant postsynaptic response (Gonzalez-Lozano et al. 2016). Like the essential post-synaptic scaffolding protein PSD-95, we observed enrichment of TRIM67 in the PSD fraction of juvenile (3-week-old) mice (**Fig 1B**). Consistent with TRIM67 expression patterns in forebrain homogenate, we saw a decrease in the abundance of TRIM67 in the PSD as mice progressed into adulthood (**Fig 1C**).

**Fig. 1.**
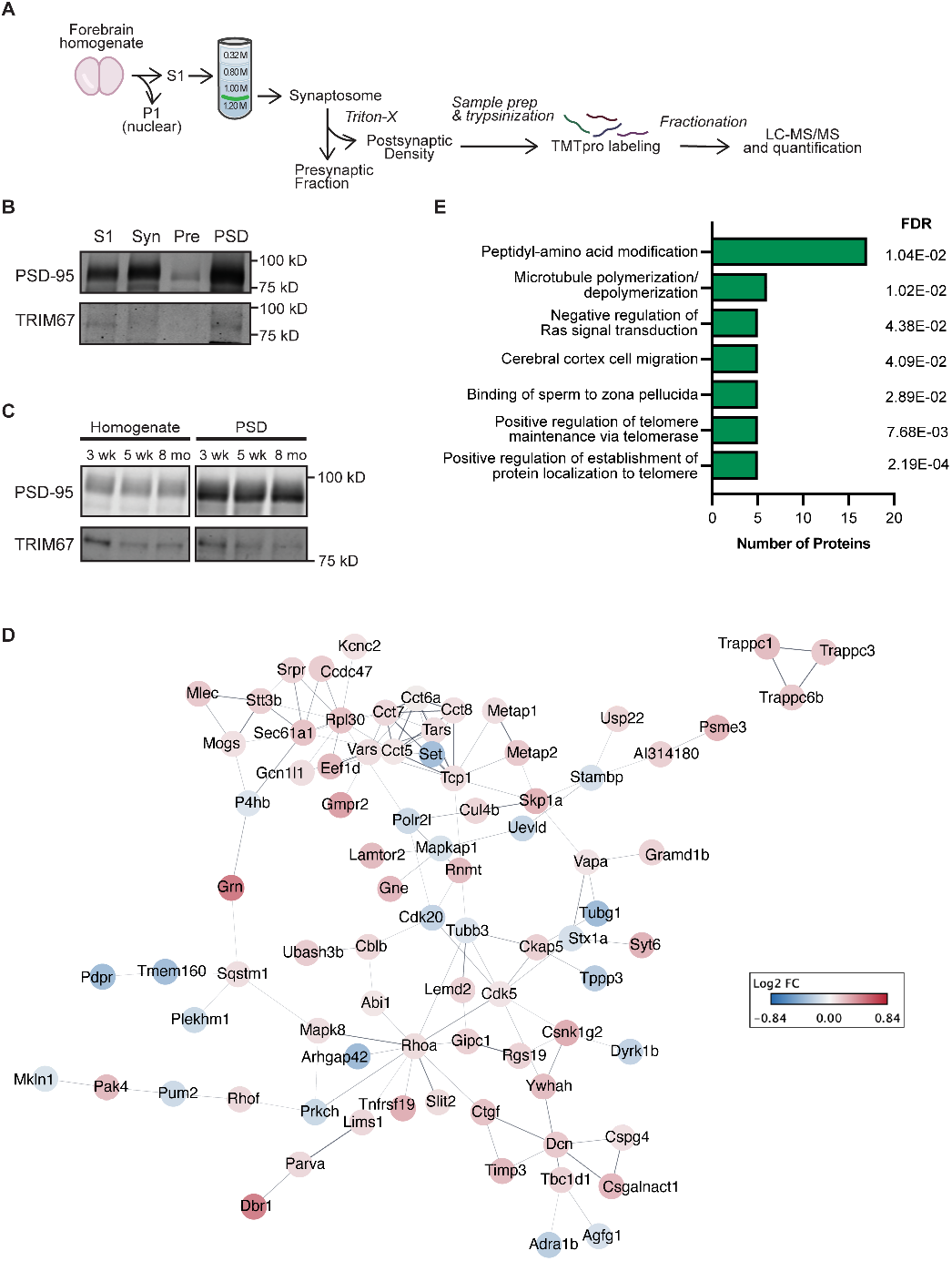
Loss of the E3 ubiquitin ligase TRIM67 alters the proteome of the post-synaptic density. **(A)** Enrichment of the post-synaptic density (PSD) fraction by differential centrifugation and downstream proteomic steps. **(B)** Immunoblotting of fractions from the PSD-enrichment of 3 week old mice. Syn: synaptosome. Pre: pre-synaptic fraction. **(C)** Immunoblotting of starting homogenate and PSD-enriched fractions from 3 week old, 5 week old, and 8 month old mice. **(D)** STRING analysis of statistically significantly changed proteins in *Trim67*^*-/-*^ mice compared to *Trim67*^*+/+*^ littermates. Lines represent published interactions between proteins. Protein color corresponds to log2 fold change of protein levels. Only proteins demonstrated to interact with at least one other significantly changed protein are visualized here. **(E)** Gene ontology analysis of significantly changed proteins by biological process.

To gain insight into the synaptic role of TRIM67, we performed proteomic analysis on the PSD-enriched fractions of *Trim67*^+/+^ and *Trim67*^-/-^ littermates. More than 7,000 proteins were detected in the PSD and we identified 149 statistically significant proteins (p <0.05) that were changed between the two genotypes (**Fig 1D**). Gene Ontology (GO) analysis of these hits demonstrated that several categories of cytoskeletal proteins were changed, including 17 proteins associated with peptidyl-amino acid modifications, six proteins involved in microtubule dynamics, and five proteins associated with negative regulation of Ras signal transduction (**Fig 1E, Extended Data**).

Although the changes in these proteins still require validation by additional methods, these data suggest that TRIM67 may influence cell signaling and cytoskeletal dynamics at the synapse. Interestingly, we observed changes in numerous proteins classified as peptidyl-amino acid modifiers (**Fig 1E, Extended Data**). This broad group includes numerous kinases and proteases, along with other protein modifying enzymes. Although no previous work has linked TRIM67 to these protein families, these proteomic results may suggest TRIM67 is required for proper signal transduction.

Furthermore, the alteration of several proteins involved in microtubule organization (**Fig 1E, Extended Data**)—including γ-tubulin (log_2_FC(−0.34))—is consistent with previously published literature on Class I TRIM proteins. The COS domain within Class I TRIM proteins mediates microtubule binding (Short and Cox 2006). In mammalian neurons, Class I TRIM46 regulates microtubules at the axon initial segment (van Beuningen et al. 2015; Harterink et al. 2019) and the single Class I TRIM ortholog in *Drosophila* helps orient new microtubule polymerization (Feng et al. 2021). As microtubules enter dendritic spines to deliver cargo in response to synaptic activity (Hu et al. 2008; McVicker et al. 2016; Schätzle et al. 2018), TRIM67 may be required for proper regulation of this polymerization in dendritic spines.

We also observed that several proteins associated with the negative regulation of Ras signal transduction were bidirectionally changed in the *Trim67*^*-/-*^ PSD (**Fig 1E, Extended Data**). Along with other small GTPases, Ras signaling is essential for the actin cytoskeleton remodeling that occurs in response to synaptic activity (Penzes and Rafalovich 2012; Zhu et al. 2002). Interestingly, previous work showed TRIM67 ubiquitinates Protein kinase C substrate 80K-H (80K-H) in the Ras pathway and reducing TRIM67 levels by siRNA increased Ras activity in a neuroblastoma cell line (Yaguchi et al. 2012). Furthermore, we observed protein level changes in five proteins associated with cerebral cortex cell migration (**Fig 1E, Extended Data**), including additional actin and microtubule regulators, as well as the morphogens that initiate their reorganization. As TRIM67 is required for morphological changes downstream of netrin-1 in the growth cone (Boyer et al. 2020), it is reasonable to hypothesize that the ligase is a necessary component for cytoskeletal signaling pathways at the synapse as well.

Lastly, we observed enrichment of five proteins involved in binding of sperm to zona pellucida and the positive regulation of telomere maintenance/protein localization to the telomere. Upon examination of the results, we discovered that these were the same five proteins—five of the eight subunits of the T-complex polypeptide-1 (TCP1) chaperone (**Fig 1E, Extended Data**). Strikingly, TCP1 serves as a multi-faceted regulator of the cytoskeleton. In addition to folding actin and /β tubulin (Sternlicht et al. 1993; Yaffe et al. 1992), TCP1 binds and inhibits the activity of gelsolin, an actin capping and severing protein (Brackley and Grantham 2011; Svanström and Grantham 2016). Furthermore, certain TCP1 subunits bind and reduce huntingtin aggregation in vitro (Behrends et al. 2006; Tam et al. 2006) and in cultured neurons (Zhao et al. 2016). In our results, we observe small (log_2_FC (0.04-0.11)) increases in TCP1 subunits. If TCP1 level changes are validated in future work, it will be imperative to determine if this is a direct effect of *Trim67* loss or a compensatory mechanism downstream of other protein level changes.

Cumulatively, our results demonstrate that TRIM67 localizes to the synapse and its loss alters the protein composition of the PSD. Of note, these studies were completed with equal numbers of male and female mice. Recently, our investigation of TRIM9 at the synapse revealed sex-dependent differences in Arp2/3 levels in *Trim9*^*-/-*^ mice (McCormick et al. 2024). Due to the high conservation between TRIM9 and TRIM67, it will be crucial to determine if TRIM67 displays these disparities as well.

## Data Availability

The proteomics dataset generated and analyzed in this study is available in the Proteomics Identification Database (PRIDE) repository (Perez-Riverol et al. 2016, 2022) under project identifier PXD048045. This dataset also includes proteomic results of *Trim9*^*+/+*^ and *Trim9*^*-/-*^ mice previously published (McCormick et al. 2024).

## Extended Data

The first sheet of the extended data spreadsheet lists significantly changed proteins in the PSD-enriched fraction of *Trim67*^-/-^ mice. For each protein, the accession code, description, gene name, coverage (%), number of peptides, number of unique peptides, number of PSM counts, log_2_fold-change, and p-value is listed. The second sheet specifically lists the proteins included in the Gene Ontology analysis results and is grouped by biological process.

## Methods

### Animals

All mouse lines were on a C57BL/6J background and bred at the University of North Carolina with approval from the Institutional Animal Care and Use Committee. Creation of the *Trim67*^-/-^ mouse line was previously described (Boyer et al. 2018).

### PSD Enrichment

Three-week old *Trim67*^*+/+*^ and *Trim67*^-/-^ littermates were identified by genotyping mice from *Trim67*^*+/-*^ breeding pairs. After sacrifice, the forebrain (cortex and hippocampus) was dissected in ice-cold dissection media, transferred to an Eppendorf and flash frozen in liquid nitrogen. Tissue was stored at –80°C until use.

Isolation of the PSD-enriched fraction was previously described (McCormick et al. 2024). Briefly, tissue was homogenized in ice cold homogenization buffer (10 mM HEPES pH 7.4, 320 mM sucrose, 1 mM EDTA, 5 mM sodium pyrophosphate, 1 mM sodium vanadate, 150 μg/mL PMSF, 2 μg/mL leupeptin, 2 μg/mL aprotinin, and 50 μM PR-619). After clearing the nuclear fraction (800 *x g*, 10 min), the total membrane fraction was pelleted (16,000 *x g*, 20 min). After resuspension in fresh homogenization buffer, the pellet was loaded onto a discontinuous sucrose gradient containing 1.2 M, 1 M or 0.8 M sucrose, as well as the inhibitors listed above. Following centrifugation in a SW-41 rotor (82.5 k *x g*, 90 min), the synaptosome fraction (the 1M/1.2M interface) was collected and pelleted at 100k *x g* (30 min). The synaptosome pellet was resuspended in 50 mM HEPES buffer with 0.5% Triton (plus inhibitors), rotated for 15 min, and centrifuged at 32k *x g* (20 min). The pellet (post-synaptic density-enriched fraction) was resuspended in 50 mM HEPES and flash frozen in LN_2_.

Following protein concentration quantification by Bradford, 100 μg of protein was aliquoted from each PSD-enriched sample. Two PSD samples of each condition (genotype x sex) were combined and a total of four replicates (from eight mice) for each condition were submitted to the UNC Hooker Proteomics Core.

### Proteomics

Proteomic methods were previously described (McCormick et al. 2024) and are reiterated below. Lysates (0.2 mg per sample; 4 replicates per condition) were precipitated using 4x cold acetone and stored at -20ºC overnight. The next day, samples were centrifuged at 15000xg at 4 ºC for 15 min, then protein pellets were reconstituted in 8M urea. All samples were reduced with 5mM DTT for 45 min at 37ºC, alkylated with 15mM iodoacetamide for 30 min in the dark at room temperature, then diluted to 1M urea with 50mM ammonium bicarbonate (pH 7.8). Samples were digested with LysC (Wako, 1:50 w/w) for 2 hr at 37ºC, then digested with trypsin (Promega, 1:50 w/w) overnight at 37ºC. The resulting peptide samples were acidified, desalted using desalting spin columns (Thermo), then the eluates were dried via vacuum centrifugation. Peptide concentration was determined using Quantitative Colorimetric Peptide Assay (Pierce).

A total of 16 samples (50 μg each) were labeled with TMT- pro reagents (Thermo Fisher) for 1 hr at room temperature. Prior to quenching, the labeling efficiency was evaluated by LC-MS/MS analysis. After confirming >98% efficiency, samples were quenched with 50% hydroxylamine to a final concentration of 0.4%. Labeled peptide samples were combined 1:1, desalted using Thermo desalting spin column, and dried via vacuum centrifugation. The dried TMT-labeled sample was fractionated using high pH reversed phase HPLC (Mertins et al. 2018). Briefly, the samples were offline fractionated over a 90 min run, into 96 fractions by high pH reverse-phase HPLC (Agilent 1260) using an Agilent Zorbax 300 Extend-C18 column (3.5-μm, 4.6 × 250 mm) with mobile phase A containing 4.5 mM ammonium formate (pH 10) in 2% (vol/vol) LC-MS grade acetonitrile, and mobile phase B containing 4.5 mM ammonium formate (pH 10) in 90% (vol/vol) LC-MS grade acetonitrile. The 96 resulting fractions were then concatenated in a non-continuous manner into 24 fractions and 5% of each were aliquoted, dried down via vacuum centrifugation and stored at -80ºC until further analysis.

The 24 TMT labeled proteome fractions were analyzed by LC/MS/MS using an Easy nLC 1200-Orbitrap Fusion Lumos (Thermo Scientific). Samples were injected onto an Easy Spray PepMap C18 column (75 μm id × 25 cm, 2 μm particle size) (Thermo Scientific) and separated over a 120 min method. The gradient for separation consisted of 5–42% mobile phase B at a 250 nl/min flow rate, where mobile phase A was 0.1% formic acid in water and mobile phase B consisted of 0.1% formic acid in 80% ACN. The Lumos was operated in SPS-MS3 mode (McAlister et al. 2014), with a 3s cycle time. Resolution for the precursor scan (m/z 400–1500) was set to 120,000 with a AGC target set to standard and a maximum injection time of 50 ms. MS2 scans consisted of CID normalized collision energy (NCE) 32; AGC target set to standard; maximum injection time of 50 ms; isolation window of 0.7 Da. Following MS2 acquisition, MS3 spectra were collected in SPS mode (10 scans per outcome); HCD set to 55; resolution set to 50,000; scan range set to 100-500; AGC target set to 200% with a 100 ms maximum inject time.

Raw data files were processed using Proteome Discoverer version 2.5, set to ‘reporter ion MS3’ with ‘16pex TMT’. Peak lists were searched against a reviewed Uniprot mouse database (downloaded Feb 2021 containing 17,051 sequences), appended with a common contaminants database, using Sequest HT within Proteome Discoverer. Data were searched with up to two missed trypsin cleavage sites, fixed modifications: TMT16plex peptide N-terminus and Lys, carbamidomethylation Cys, dynamic modification: N-terminal protein acetyl, oxidation Met. Precursor mass tolerance of 10ppm and fragment mass tolerance of 0.5 Da (MS3). Peptide false discovery rate was set to 1%. Reporter abundance was calculated based on intensity; for MS3 data, SPS mass matches threshold was set to 50 and co-isolation threshold was set to 100. Razor and unique peptides were used for quantitation. Proteins with >50% missing TMT intensities across samples were removed. Student’s t-tests were conducted within Proteome Discoverer, and a p-value <0.05 was considered significant. Log_2_fold change ratios were calculated for each pairwise comparison.

Protein-protein interactions were visualized with Cytoscape (Shannon et al. 2003). Enrichment of biological processes in significantly changed proteins was identified using Gene Ontology analysis (Ashburner et al. 2000; The Gene Ontology Consortium et al. 2023; Thomas et al. 2022).

## ACKNOWLEDGEMENTS

We thank Vong Thoong, Christopher Hardie, Mayra Correa-Ramirez, and Janee Cadlett-Jette for assistance with mouse husbandry and genotyping. We thank Graham Diering for experimental design advice and PSD isolation guidance.

This work was supported by National Institutes of Health Grants R35GM135160 (S.L.G.) and 1F31NS113381-01 (L.E.M). Mass spectrometry was performed at the UNC Proteomics Core Facility, which is supported in part by NCI Center Core Support Grant (2P30CA016086-45) to the UNC Lineberger Comprehensive Cancer Center. Mass spectrometry was also supported by an award from the UNC Core Facilities Advocacy Committee and Office of Research Technologies, UNC Chapel Hill School of Medicine.

## References

Ashburner, M., Ball, C. A., Blake, J. A., Botstein, D., Butler, H., Cherry, J. M., Davis, A. P., Dolinski, K., Dwight, S. S., Eppig, J. T., Harris, M. A., Hill, D. P., Issel-Tarver, L., Kasarskis, A., Lewis, S., Matese, J. C., Richardson, J. E., Ringwald, M., Rubin, G. M., Sherlock, G. (2000). Gene Ontology: Tool for the unification of biology. Nature Genetics, 25(1), Article 1. 10.1038/75556

Behrends, C., Langer, C. A., Boteva, R., Böttcher, U. M., Stemp, M. J., Schaffar, G., Rao, B. V., Giese, A., Kretzschmar, H., Siegers, K., Hartl, F. U. (2006). Chaperonin TRiC Promotes the Assembly of polyQ Expansion Proteins into Nontoxic Oligomers. Molecular Cell, 23(6), 887–897. 10.1016/j.molcel.2006.08.017

Boyer, N. P., McCormick, L. E., Menon, S., Urbina, F. L., Gupton, S. L. (2020). A pair of E3 ubiquitin ligases compete to regulate filopodial dynamics and axon guidance. Journal of Cell Biology, 219(1), e201902088. 10.1083/jcb.201902088

Boyer, N. P., Monkiewicz, C., Menon, S., Moy, S. S., Gupton, S. L. (2018). Mammalian TRIM67 functions in brain development and behavior. eNeuro, 5(3). 10.1523/ENEURO.0186-18.2018

Brackley, K. I., Grantham, J. (2011). Interactions between the actin filament capping and severing protein gelsolin and the molecular chaperone CCT: Evidence for nonclassical substrate interactions. Cell Stress Chaperones, 16(2), 173–179. 10.1007/s12192-010-0230-x

Feng, C., Cleary, J. M., Kothe, G. O., Stone, M. C., Weiner, A. T., Hertzler, J. I., Hancock, W. O., Rolls, M. M. (2021). Trim9 and Klp61F promote polymerization of new dendritic microtubules along parallel microtubules. Journal of Cell Science, 134(11), jcs258437. 10.1242/jcs.258437

Gonzalez-Lozano, M. A., Klemmer, P., Gebuis, T., Hassan, C., van Nierop, P., van Kesteren, R. E., Smit, A. B., Li, K. W. (2016). Dynamics of the mouse brain cortical synaptic proteome during postnatal brain development. Scientific Reports, 6(1), 35456. 10.1038/srep35456

Harterink, M., Vocking, K., Pan, X., Soriano Jerez, E. M., Slenders, L., Fréal, A., Tas, R. P., van de Wetering, W. J., Timmer, K., Motshagen, J., van Beuningen, S. F. B., Kapitein, L. C., Geerts, W. J. C., Post, J. A., Hoogenraad, C. C. (2019). TRIM46 Organizes Microtubule Fasciculation in the Axon Initial Segment. The Journal of Neuroscience: The Official Journal of the Society for Neuroscience, 39(25), 4864–4873. 10.1523/JNEUROSCI.3105-18.2019

Hu, X., Viesselmann, C., Nam, S., Merriam, E., Dent, E. W. (2008). Activity-Dependent Dynamic Microtubule Invasion of Dendritic Spines. Journal of Neuroscience, 28(49), 13094–13105. 10.1523/JNEUROSCI.3074-08.2008

McAlister, G. C., Nusinow, D. P., Jedrychowski, M. P., Wühr, M., Huttlin, E. L., Erickson, B. K., Rad, R., Haas, W., Gygi, S. P. (2014). MultiNotch MS3 enables accurate, sensitive, and multiplexed detection of differential expression across cancer cell line proteomes. Analytical Chemistry, 86(14), 7150–7158. 10.1021/ac502040v

McCormick, L. E., Evans, E. B., Barker, N. K., Herring, L. E., Diering, G. H., Gupton, S. L. (2024). The E3 ubiquitin ligase TRIM9 regulates synaptic function and actin dynamics (p. 2023.12.31.573790). bioRxiv. 10.1101/2023.12.31.573790

McVicker, D. P., Awe, A. M., Richters, K. E., Wilson, R. L., Cowdrey, D. A., Hu, X., Chapman, E. R., Dent, E. W. (2016). Transport of a kinesin-cargo pair along microtubules into dendritic spines undergoing synaptic plasticity. Nature Communications, 7, 12741. 10.1038/ncomms12741

Menon, S., Goldfarb, D., Ho, C. T., Cloer, E. W., Boyer, N. P., Hardie, C., Bock, A. J., Johnson, E. C., Anil, J., Major, M. B., Gupton, S. L. (2021). The TRIM9/TRIM67 neuronal interactome reveals novel activators of morphogenesis. Molecular Biology of the Cell, 32(4), 314–330. 10.1091/mbc.E20-10-0622

Mertins, P., Tang, L. C., Krug, K., Clark, D. J., Gritsenko, M. A., Chen, L., Clauser, K. R., Clauss, T. R., Shah, P., Gillette, M. A., Petyuk, V. A., Thomas, S. N., Mani, D. R., Mundt, F., Moore, R. J., Hu, Y., Zhao, R., Schnaubelt, M., Keshishian, H., … Carr, S. A. (2018). Reproducible workflow for multiplexed deep-scale proteome and phosphoproteome analysis of tumor tissues by liquid chromatography–mass spectrometry. Nature Protocols, 13(7), Article 7. 10.1038/s41596-018-0006-9

Penzes, P., Rafalovich, I. (2012). Regulation of the actin cytoskeleton in dendritic spines. Advances in Experimental Medicine and Biology, 970, 81–95. 10.1007/978-3-7091-0932-84

Perez-Riverol, Y., Bai, J., Bandla, C., García-Seisdedos, D., Hewapathirana, S., Kamatchinathan, S., Kundu, D. J., Prakash, A., Frericks-Zipper, A., Eisenacher, M., Walzer, M., Wang, S., Brazma, A., Vizcaíno, J. A. (2022). The PRIDE database resources in 2022: A hub for mass spectrometry-based proteomics evidences. Nucleic Acids Research, 50(D1), D543–D552. 10.1093/nar/gkab1038

Perez-Riverol, Y., Xu, Q.-W., Wang, R., Uszkoreit, J., Griss, J., Sanchez, A., Reisinger, F., Csordas, A., Ternent, T., Del-Toro, N., Dianes, J. A., Eisenacher, M., Hermjakob, H., Vizcaíno, J. A. (2016). PRIDE Inspector Toolsuite: Moving Toward a Universal Visualization Tool for Proteomics Data Standard Formats and Quality Assessment of ProteomeXchange Datasets. Molecular Cellular Proteomics: MCP, 15(1), 305–317. 10.1074/mcp.O115.050229

Schätzle, P., Esteves Da Silva, M., Tas, R. P., Katrukha, E. A., Hu, H. Y., Wierenga, C. J., Kapitein, L. C., Hoogenraad, C. C. (2018). Activity-Dependent Actin Remodeling at the Base of Dendritic Spines Promotes Microtubule Entry. Current Biology, 28(13), 2081–2093.e6. 10.1016/j.cub.2018.05.004

Shannon, P., Markiel, A., Ozier, O., Baliga, N. S., Wang, J. T., Ramage, D., Amin, N., Schwikowski, B., Ideker, T. (2003). Cytoscape: A Software Environment for Integrated Models of Biomolecular Interaction Networks. Genome Research, 13(11), 2498–2504. 10.1101/gr.1239303

Short, K. M., Cox, T. C. (2006). Subclassification of the RBCC/TRIM Superfamily Reveals a Novel Motif Necessary for Microtubule Binding *. Journal of Biological Chemistry, 281(13), 8970–8980. 10.1074/jbc.M512755200

Sternlicht, H., Farr, G. W., Sternlicht, M. L., Driscoll, J. K., Willison, K., Yaffe, M. B. (1993). The t-complex polypeptide 1 complex is a chaperonin for tubulin and actin in vivo. Proceedings of the National Academy of Sciences of the United States of America, 90(20), 9422–9426.

Svanström, A., Grantham, J. (2016). The molecular chaperone CCT modulates the activity of the actin filament severing and capping protein gelsolin in vitro. Cell Stress Chaperones, 21(1), 55–62. 10.1007/s12192-015-0637-5

Tam, S., Geller, R., Spiess, C., Frydman, J. (2006). The chaperonin TRiC controls polyglutamine aggregation and toxicity through subunit-specific interactions. Nature Cell Biology, 8(10), Article 10. 10.1038/ncb1477

The Gene Ontology Consortium, Aleksander, S. A., Balhoff, J., Carbon, S., Cherry, J. M., Drabkin, H. J., Ebert, D., Feuermann, M., Gaudet, P., Harris, N. L., Hill, D. P., Lee, R., Mi, H., Moxon, S., Mungall, C. J., Muruganugan, A., Mushayahama, T., Sternberg, P. W., Thomas, P. D., … Westerfield, M. (2023). The Gene Ontology knowledgebase in 2023. Genetics, 224(1), iyad031. 10.1093/genetics/iyad031

Thomas, P. D., Ebert, D., Muruganujan, A., Mushayahama, T., Albou, L.-P., Mi, H. (2022). PANTHER: Making genome-scale phylogenetics accessible to all. Protein Science, 31(1), 8–22. 10.1002/pro.4218

van Beuningen, S. F. B., Will, L., Harterink, M., Chazeau, A., van Battum, E. Y., Frias, C. P., Franker, M. A. M., Katrukha, E. A., Stucchi, R., Vocking, K., Antunes, A. T., Slenders, L., Doulkeridou, S., Sillevis Smitt, P., Altelaar, A. F. M., Post, J. A., Akhmanova, A., Pasterkamp, R. J., Kapitein, L. C., … Hoogenraad, C. C. (2015). TRIM46 Controls Neuronal Polarity and Axon Specification by Driving the Formation of Parallel Microtubule Arrays. Neuron, 88(6), 1208–1226. 10.1016/j.neuron.2015.11.012

Yaffe, M. B., Farr, G. W., Miklos, D., Horwich, A. L., Sternlicht, M. L., Sternlicht, H. (1992). TCP1 complex is a molecular chaperone in tubulin biogenesis. Nature, 358(6383), 245–248. 10.1038/358245a0

Yaguchi, H., Okumura, F., Takahashi, H., Kano, T., Kameda, H., Uchigashima, M., Tanaka, S., Watanabe, M., Sasaki, H., Hatakeyama, S. (2012). TRIM67 Protein Negatively Regulates Ras Activity through Degradation of 80K-H and Induces Neuritogenesis. The Journal of Biological Chemistry, 287(15), 12050–12059. 10.1074/jbc.M111.307678

Zhao, X., Chen, X.-Q., Han, E., Hu, Y., Paik, P., Ding, Z., Overman, J., Lau, A. L., Shahmoradian, S. H., Chiu, W., Thompson, L. M., Wu, C., Mobley, W. C. (2016). TRiC subunits enhance BDNF axonal transport and rescue striatal atrophy in Huntington’s disease. Proceedings of the National Academy of Sciences of the United States of America, 113(38), E5655–5664. 10.1073/pnas.1603020113

Zhu, J. J., Qin, Y., Zhao, M., Van Aelst, L., Malinow, R. (2002). Ras and Rap control AMPA receptor trafficking during synaptic plasticity. Cell, 110(4), 443–455. 10.1016/s0092-8674(02)00897-8

